# CIB2 regulates autophagy via Rheb-mTORC1 signaling axis

**DOI:** 10.1101/2020.09.18.302265

**Authors:** Saumil Sethna, Steven L. Bernstein, Xiaoying Jian, Sheikh Riazuddin, Paul A. Randazzo, Saima Riazuddin, Zubair M. Ahmed

## Abstract

Age-related macular degeneration (AMD), a multifactorial neurodegenerative disorder, is the most common cause of vision loss in the elderly. Deficits in autophagy have been associated with age-related retinal pigment epithelium (RPE) pathology in mice, and dry-AMD in humans. In this study, we establish that the calcium and integrin binding protein 2 (CIB2) regulates autophagy in the RPE via Rheb-mTORC1 signaling axis. *Cib2* mutant mice have reduced autophagic clearance in RPE and increased mTORC1 signaling – a negative regulator of autophagy. Concordant molecular deficits were also observed in RPE/choroid tissues from humans affected with dry AMD. Mechanistically, CIB2 negatively regulates mTORC1 by preferentially binding to ‘nucleotide empty’ or inactive GDP-loaded Rheb. Upregulated mTORC1 signaling has been implicated in aging, Tuberous sclerosis complex (TSC), and lymphangioleiomyomatosis (LAM) cancer. Over-expressing CIB2 in LAM patient-derived fibroblasts and *Tsc2* null cell line down-regulates hyperactive mTORC1 signaling. Thus, our findings have significant ramifications for the etiology of AMD and mTORC1 hyperactivity disorders and treatments.

## INTRODUCTION

Age-related macular degeneration (AMD) is a progressive degenerative disease of the macula, a specialized region of the retina responsible for daytime vision. AMD often causes central vision loss and irreversible blindness, and affects 10% of the population aged 65-75 years and 25% of those aged ≥75 years. By 2050, its prevalence is expected to increase by 50% ^1,2^. AMD is categorized as either ‘wet’ (associated with choroidal neovascularization) or ‘dry’ (associated with atrophy and progressive thinning of retinal layers). Dry AMD occurs in ~90% of the cases and can cause permanent vision loss if left untreated ^3^. Around 35 genes have been associated with AMD and some patients with dry AMD benefit from lifestyle modifications (particularly cessation of smoking ^4–13^ or specific vitamin supplementation ^14^), however, no current therapeutic strategy actually prevents dry AMD progression.

Etiologically, dry AMD originates primarily in the retinal pigment epithelium (RPE) and choroid, and secondarily, it impairs photoreceptor (PR) function and integrity. RPE supports the PR and hence vision by fulfilling several critical functions, including non-canonical autophagy or LC3-associated phagocytosis (LAP) of PR outer segments (OS) ^15,16^. Likewise, macroautophagy (henceforth autophagy) is a catabolic process which removes cellular debris, damaged/aged organelles, and shares many features with LAP. Autophagy is an essential process for all cells, but its function is especially notable in post-mitotic cells such as those in the RPE ^16^ that have the highest life-long phagocytic load of any cell-type in the body. Defects in autophagy has been implicated in dry AMD ^17,18^. Besides AMD, deficits in autophagy are linked to lifespan and several diseases such as Alzheimer’s, Parkinson’s, certain cancers, and metabolic disorders amongst others ^19–21^. Therefore, deciphering the molecular regulators of autophagy has significant ramifications for fundamental cellular aging process and the etiology of autophagy-related disorders.

Mechanistic target of rapamycin (mTOR) is a serine/threonine kinase. mTOR forms two large multi-subunit complexes: mTORC1 and mTORC2. mTORC1 is a negative regulator of autophagy, and it also integrates nutrient and growth factors via its nutrient sensing arm and tuberous sclerosis complex (TSC) complex, respectively ^22,23^. TSC complex (consists of TSC1, TSC2, and TBC1D7) directly regulates the small GTPase, Rheb (Ras homolog enriched in brain) via spatial control and TSC2’s Rheb-GAP activity ^24^. Rheb associates with lysosomal surface through its C-terminus farnesyl group ^25^. GTP bound Rheb is a potent and obligate activator of mTORC1 that acts via allosterically realigning the kinase active site of mTORC1 ^26^. Aberrant mTORC1 signaling is implicated in ageing, cancers, obesity, and Alzheimer’s amongst others ^27^. Therefore, identifying modulators of mTORC1 pathway has significant basic cellular and clinical implications. Here, we report that RPE-specific ablation of CIB2 leads to dysregulation of mTORC1 and autophagy. Our biochemical and interaction studies reveal the preferential binding of CIB2 to the inactive state of Rheb, which then acts as a negative regulator of mTORC1.

## RESULTS

### Loss of CIB2 leads to autophagic clearance defects due to hyperactive mTORC1 signaling

In the parallel study, we showed that removal of single or both copies of *Cib2,* specifically in the RPE, leads to age-associated phenotypes including sub-RPE deposits, accumulation of proteins and lipid species found in drusen, and reduced electroretinogram (ERG) amplitudes (**Fig. 1a**) ^28^. Lack of CIB2 leads to defects in LC3-associated phagocytosis and reduced levels of autophagy proteins such as ATG5-12, in the RPE ^28^. Non-canonical autophagy such as phagolysosomal OS processing shares many aspects and proteins with canonical autophagy ^16^. Hence, we reasoned that autophagy is impaired in *Cib2* mutant mice. When autophagy is fully functional, the levels of autophagosome membrane protein LC3-II increase, while those of the autophagosome marker protein p62/SQSTM1 decrease in concert, due to specific digestion within the autolysosome. When autophagy is faulty, levels of both p62/SQSTM1 and LC3-II increase ^29^. Thus, to assess the functionality, we evaluated the levels of p62/SQSTM1 and LC3-II in *Cib2* mutant mice after inducing autophagy.

**Figure 1:**
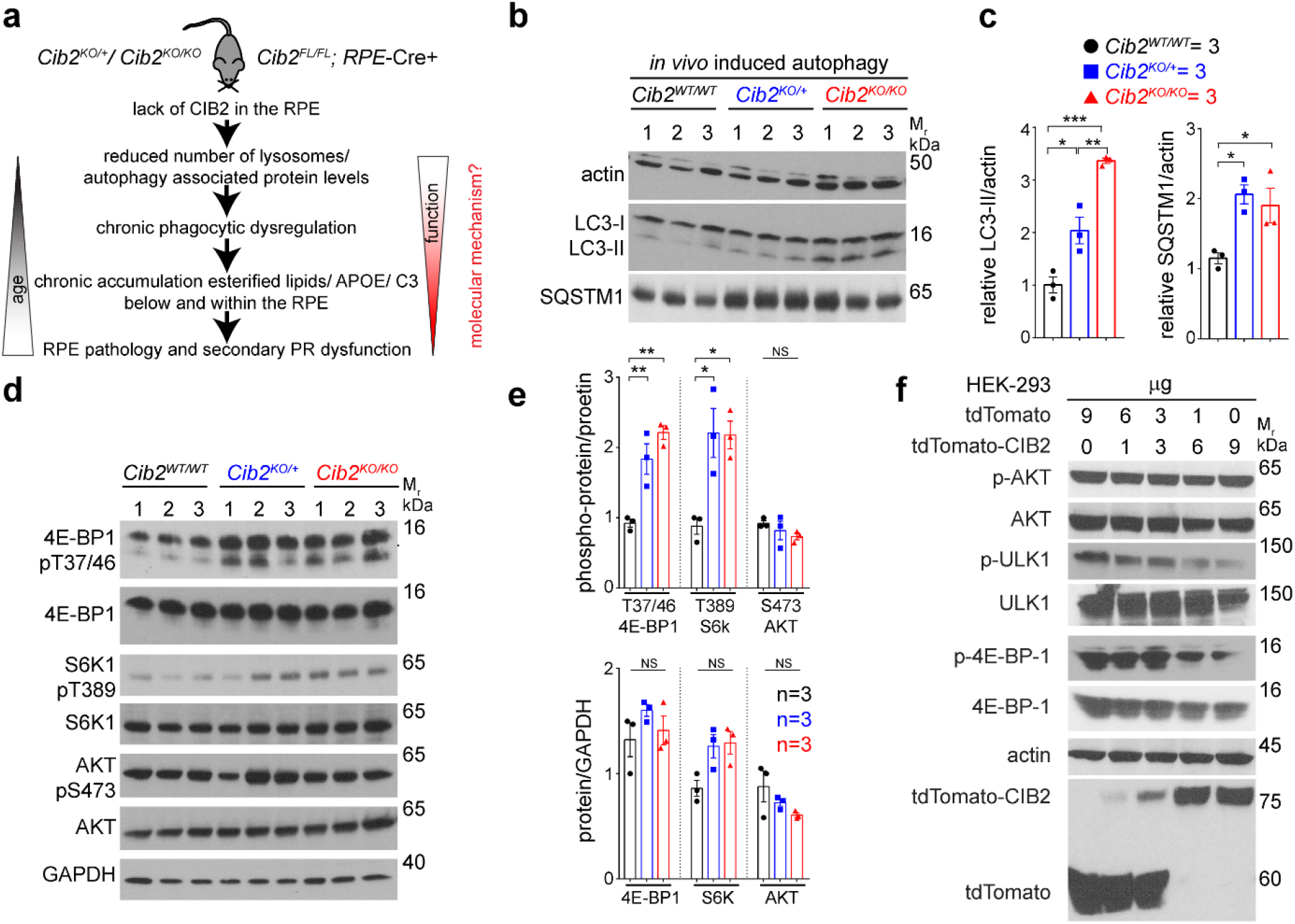
CIB2 modulates autophagy via mTORC1 pathway. **a** Summary of eye disorder found in *Cib2* mutant mice. Global loss of single or both alleles of *Cib2* (*Cib2^KO/+^, Cib2^KO/KO^*), or specifically within the RPE (*Cib2^FL/FL^.* RPE-*Cre+*) lead to reduced non-canonical autophagy and chronic phagolysosomal processing defects, which gets exacerbated with age leading to lipid and canonical AMD drusen markers accumulation below the RPE and secondarily leads to age-associated loss of photoreceptor (PR) function. **b, c** Induced autophagy *in vivo* by food starving for 24 hrs shows dysregulated autophagy in RPE/choroid lysates, as levels of LC3-II and p62/SQSTM1 are higher, quantified in panel **b**, in *Cib2* mutant mice. **d, e** Higher levels, quantified in panel **e**, of phospho-protein levels of mTORC1 targets 4-EB-P1 and S6K1, but not mTORC2 target AKT2, indicate hyperactive mTORC1 in RPE/choroid lysates after *in vivo* induced autophagy. **f** Representative immunoblots of indicated phosphoproteins and proteins from HEK293 lysates over-expressing increasing amounts of tdTomato-CIB2 and decreasing amounts of tdTomato (total amount of plasmid DNA for each condition is 9 μg) causing reduction in mTORC1 activity. One-way ANOVA and Bonferroni *post hoc* test (**c, e**). *p* value of <0.05 (*), <0.01 (**), or <0.001 (***), NS – not significant.

First, to induce autophagy *in vivo,* we fasted 2-3-month-old animals for 24 hrs and collected the RPE/choroid. Immunoblots of RPE/choroid lysates from autophagy induced *Cib2^KO/+^* and *Cib2^KO/KO^* mice showed that their p62/SQSTM1 and LC3-II levels doubled compared to those in WT age-matched controls (**Fig. 1b, c**). To assess if autophagy is dysregulated over time, we assessed the LC3-II and p62/SQSTM1 levels in aged mice, fed ad-libitum. We observed 2-3-fold increase in both LC3-II and p62/SQSTM1 levels, which suggests that loss of CIB2 leads to early as well as age-related autophagy deficits (**supplementary Fig. 1a, b**). LC3-I is a cytosolic protein but when lipidated with a phosphatidylethanolamine adduct (LC3-II form) localizes to the growing auto/phagosome membrane, and thus can be used as a reliable readout for the dynamic autophagic process ^16,29,30^. RPE/choroid whole mounts, *ex vivo*, were incubated overnight with DMSO (control) or bafilomycin A1, which blocks autophagosome fusion with lysosomes, and the relative ratio of LC3-II (bafilomycin A1/DMSO) was assessed with immunoblotting ^29^. We found ~50% reduction in relative LC3 flux in RPE-specific *Cib2^KO^* mice (**supplementary Fig. 1c, d**). In addition, overexpressing CIB2 in RPE-J cells markedly increased relative LC3 flux in the presence of bafilomycin A1. The mTORC1 inhibitor rapamycin similarly elevated LC3-II levels (**supplementary Fig. 1e, f**). These results strongly support the role of CIB2 in clearance of autophagosomes, without compromising autophagosome biogenesis.

Since mTORC1 is a known negative regulator of autophagy, we assessed the status of mTORC1 signaling in *Cib2* mutant mice. First, we measured phosphoprotein levels of the bonafide mTORC1 downstream targets 4E binding protein 1 (4E-BP1) and S6 kinase 1 (S6K1). RPE/choroid lysate immunoblots from mutant mice undergoing induced autophagy revealed an approximate 2-fold increase in 4E-BP1 and S6K1 phospho-protein levels, but we found no difference in levels of the mTORC2 downstream target phospho-AKT. (**Fig. 1d, e**). mTORC1 integrates various metabolic and amino acid signals through its nutrient-sensing arm and growth signals, which converge on the small GTPase Rheb, via the TSC complex ^23^. Next, we tested if CIB2 is a specific negative regulator of mTORC1, mTORC2 or both. For this purpose, we used HEK293 cells, because the mTORC1 signaling pathway is well-established in these cells ^31^, and also to expand CIB2 functions beyond the RPE. HEK293 cells overexpressing increasing amounts of CIB2 showed decreasing phosphorylation of the mTORC1 targets 4E-BP1, and ULK1. We found no changes in mTORC2-mediated phosphorylation of AKT (**Fig. 1f**). Together, the results suggest that CIB2 impacts autophagy, specifically by modulating mTORC1.

### CIB2 forms a tripartite complex with Raptor-mTOR

To further explore the molecular mechanism of CIB2-mediated mTORC1 modulation, we assessed direct interactions between CIB2 and TSC1, TSC2, mTORC1-specifc subunit Raptor, mTOR kinase, and Rheb, using a previously reported nanoscale pulldown 2.0 (NanoSPD) quantitative interaction assay ^32^. NanoSPD utilizes a construct (nanoTRAP) consisting of GFP nanobody fused with the heavy meromyosin (HMM domain) of myosin 10 (myo10^HMM^-GFP nanobody). This allows preferential migration of any GFP-tagged protein to the filopodia tips in the transfected cells ^32^. As anticipated, GFP-hCIB2 (bait) migrated efficiently to filopodia in COS-7 cells only in the presence of the nanoTRAP. As control, we ruled out the interaction of HA-mCherry-Raptor or myc-mTOR with nanoTRAP and with GFP tag (**Fig. 2a, d and data not shown**). Using NanoSPD assay, TSC1 and TSC2, the major proteins of the TSC complex directly regulating Rheb, did not show any co-accumulation with GFP-hCIB2 (**Fig. 2c, d**). In contrast, we found that Raptor, but not mTOR, interacts with CIB2. However, mTOR and CIB2 form a tripartite complex in the presence of Raptor (**Fig. 2b, d**). We further confirmed the CIB2-Raptor interaction using co-immunoprecipitation (co-IP) assay (**Fig. 2e**).

**Figure 2:**
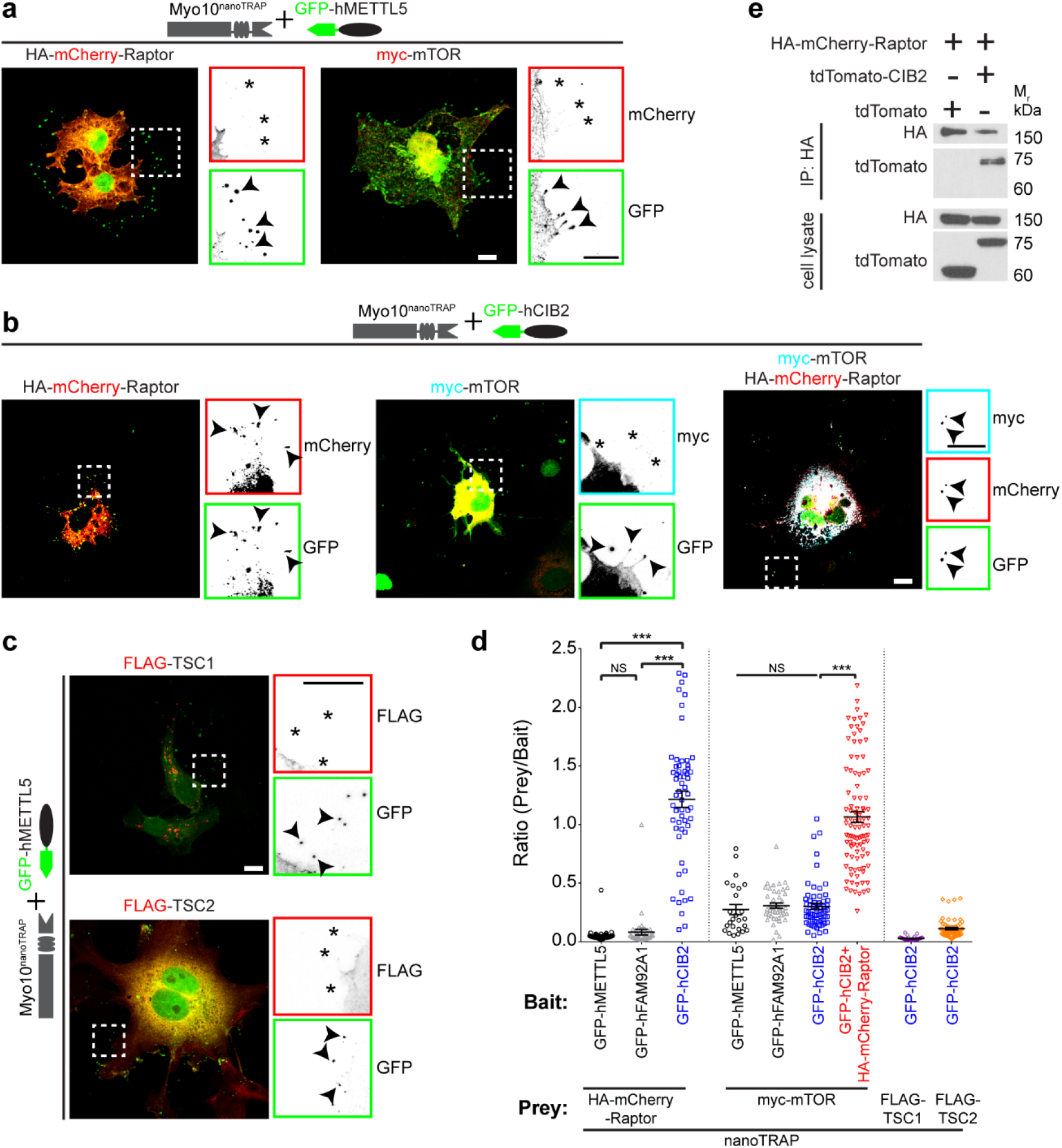
CIB2 interacts with Raptor and forms a tripartite complex with mTOR via Raptor. **a** Merged confocal micrographs of the HA-mCherry-Raptor (*left* panel) and myc-mTOR (*right* panel) constructs transiently transfected with nanoTRAP and GFP-hMETTL5 (controls) in COS-7 cells. Zoomed images of the inset shows reverse color images of indicated constructs at the tip of filopodia. *Arrowheads* indicate accumulation of GFP tagged protein at the tip of filopodia, * indicates absence of the indicated construct in the red channel at the filopodia tips, suggesting no interaction of tagged proteins with the nanoTRAP and control GFP-METTL5 construct. Similar results were obtained with nanoTRAP and GFP-FAM92A1 (control) construct. **b** Merged images for indicated constructs in *green:* GFP-hCIB2, *cyan:* myc-mTOR, *red*: HA-mCherry-Raptor indicating that mTOR does not interact directly with CIB2 but forms a tripartite complex with Raptor and CIB2. Zoomed images of the inset shows reverse color images of indicated constructs at the tip of filopodia. **c** Confocal images for FLAG-tagged TSC1 (*top* panel) and TSC2 (*bottom* panel) and nanoTRAP plus GFP-hCIB2 shows no interaction for either protein with hCIB2-eGFP. **d** Quantification for nanoTRAP assay for images shown in panels **a-c**. *** *p*< 0.001 by one-way ANOVA. **e** Co-IP experiments in HEK293 cells show interaction between exogenously expressed Raptor and CIB2. *Scale bar:* 10 μm.

mTORC1 is a cytosolic complex when starved of amino acids and nutrients. However, it localizes to the lysosomes in the presence of amino acid, nutrients, and growth factors ^22,23^. We sought to assess whether the loss of CIB2 leads to aberrant localization of mTORC1 complex. First, we generated adipose tissue-derived mesenchymal stem cells (MSCs) from WT and *Cib2KO/KO* mice (**supplementary Fig. 2**) to monitor the localization of mTORC1 complex. In both WT and *Cib2^KO/KO^* MSCs mTOR was cytosolic in the absence of amino acids, (**Fig. 3a**). Next, we triggered lysosomal localization by adding amino acids in the media. In both WT and *Cib2^KO/KO^* MSCs mTOR was co-localized with lysosomal marker LAMP, indicating that the loss of CIB2 did not affect the lysosomal targeting of mTOR (**Fig. 3a**). Similarly, we observed no localization deficits in the absence and presence of other signaling factors such as insulin (**Fig. 3b**). In WT MSCs, we also assessed the co-location of CIB2 with TSC2. We found no significant co-localization between CIB2 and TSC2 under any of the conditions tested (supplementary Fig. 3a). These findings imply that CIB2 function is beyond controlling mTORC1 complex localization to lysosomes and downstream of TSC complex.

**Figure 3:**
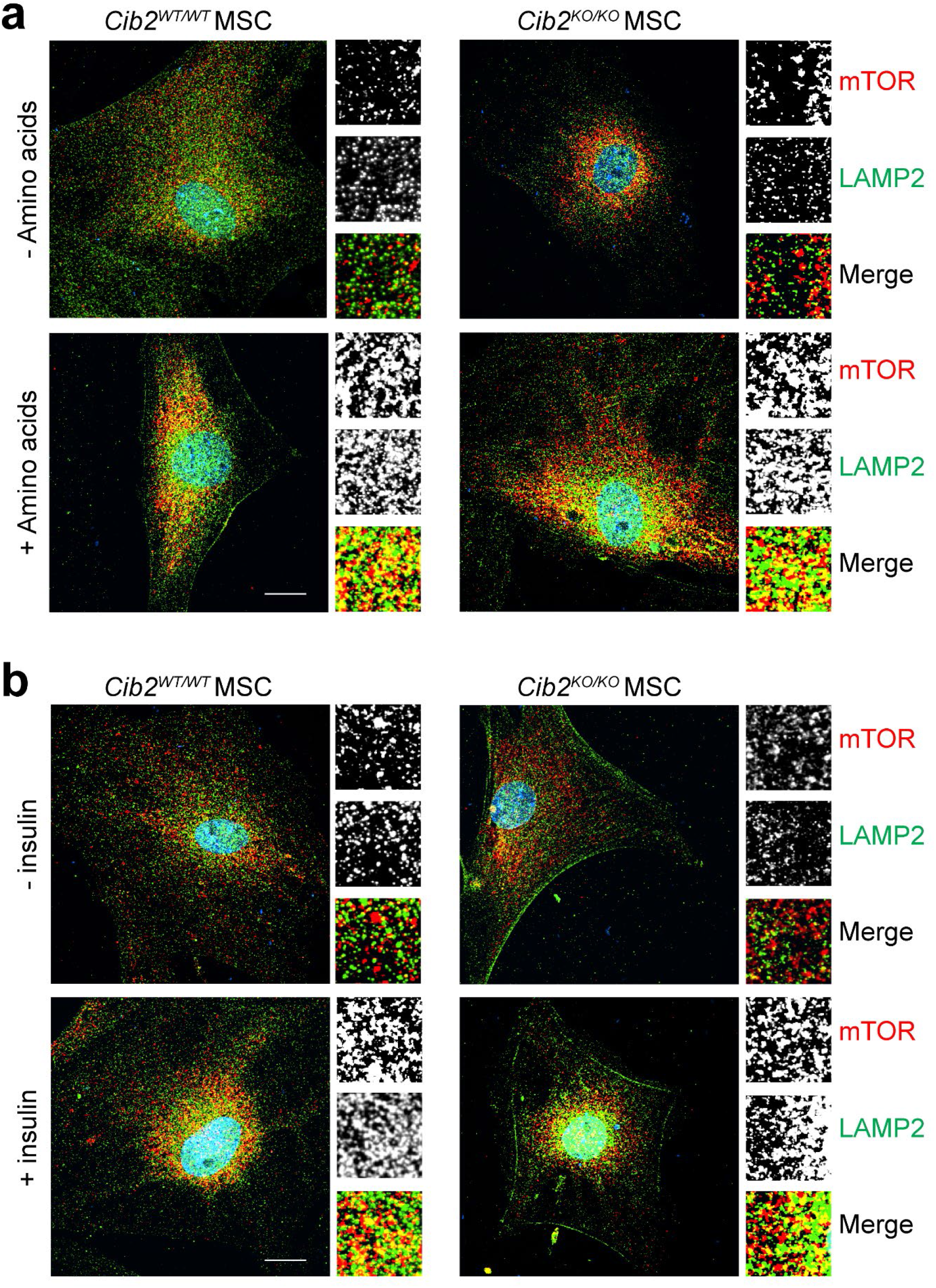
Loss of CIB2 has no apparent impact on the targeting of mTORC1 to lysosomes. **a, b** Mesenchymal stem cells (MSC) from WT (*left* panels) or *Cib2* mutant mice (*Cib2^KO/KO^)* were serum starved for 16 hrs, followed by amino acid starvation for 40 mins and stimulated with amino acids for 20 mins (+amino acids) or left unstimulated (-amino acids), or serum starved for 40 mins and stimulated with 10 nM insulin (+insulin) or treated with DMEM (-insulin). Cells were fixed and stained for lysosomal protein LAMP2 (green) or mTOR (red). Amino acid stimulation lead to co-lococalization of mTOR with lysosomes (yellow puncta) irrespective of genotype **a**. Insulin stimulation leads to partial co-localization of mTOR with lysosomes **b**. Zoomed in single color images of denoted proteins or merge. *Scale bar:* 20 μm.

### CIB2 negatively regulates mTORC1 via preferential binding to ‘nucleotide empty-’ or GDP-Rheb

When we analyzed the co-localization of CIB2 with Rheb, we found significant overlap under all the conditions tested (supplementary Fig. 3b). To assess whether CIB2 actually directly interacts with Rheb we utilized the NanoSPD assay. Interestingly, NanoSPD revealed direct interaction between CIB2 and the small GTPase Rheb (**Fig. 4a, b**), adding a new binding partner into the growing list of Rheb interactome ^33^. CIB2 partnered uniquely with Rheb (but not with other Rheb-activators), including TSC complex and microspherule protein 1 (MCRS1; **Supplementary Fig. 4a-c**). Rheb is a 184 amino acid evolutionary conserved GTPase (**supplementary Fig. 5a**). GTP bound Rheb acts as a potent and obligate activator of mTORC1 ^26^. Hence, we next explored the impact of the nucleotide binding state of Rheb on interaction with CIB2 through co-IP studies (**Fig. 4c-f**). For these studies, we used WT protein, Rheb harboring the Q64L variant that has increased GTP loading and partial resistance to GAP activity of TSC, and Rheb with S20N missense variant that has severely reduced GTP and GDP binding capacity (~1% of WT Rheb), essentially acting as a ‘nucleotide empty’ variant ^34^. Consistent with prior findings, we found reduced expression of Rheb-S20N variant protein (**Fig. 4d** – cell lysate, **supplementary Fig. 5d**). In co-IP assay, CIB2 bound preferentially to the S20N variant as compared to either the Rheb-WT or the Rheb-Q64L variant (**Fig. 4d**).

**Figure 4:**
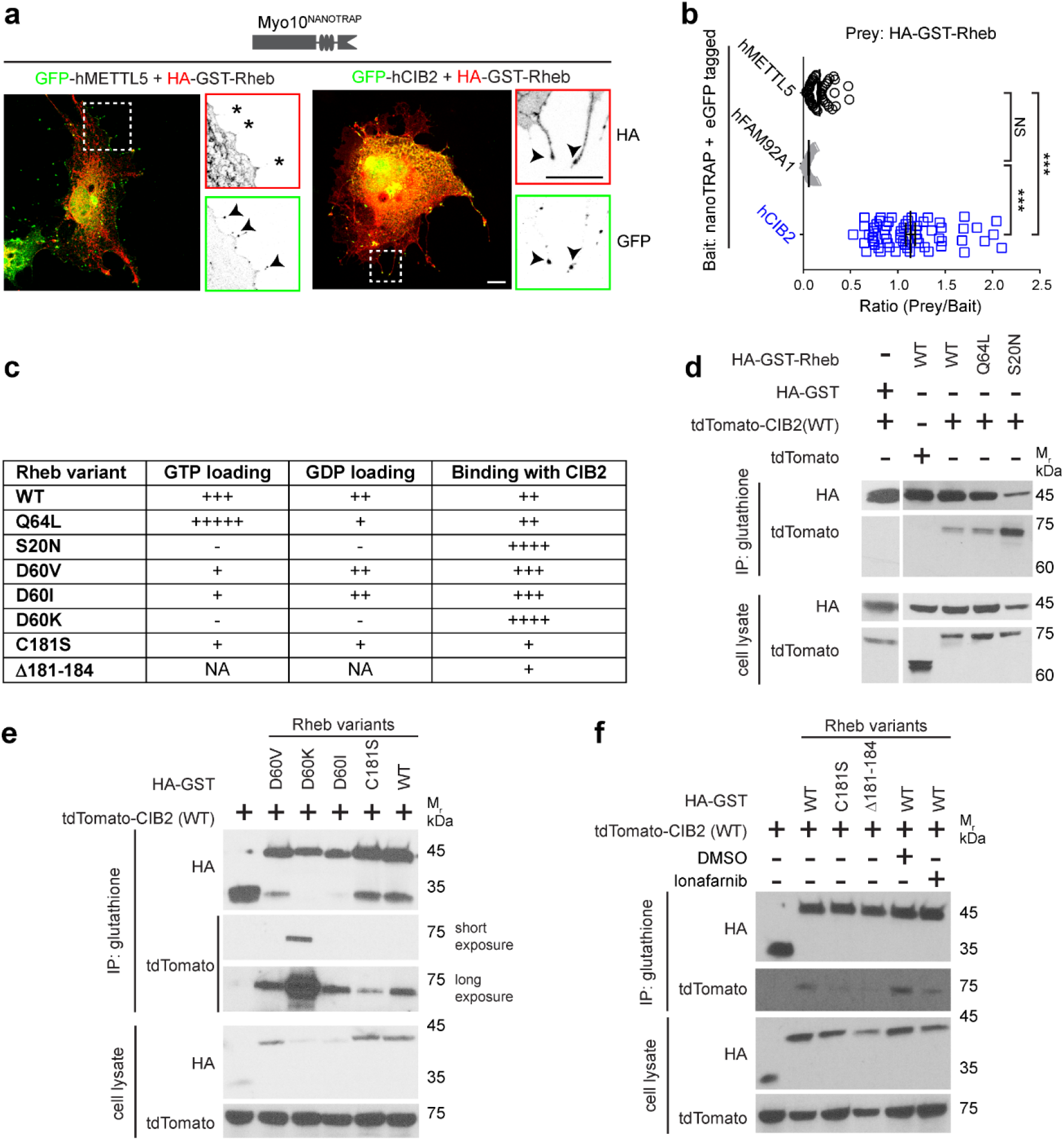
CIB2 modulates mTORC1 signaling via preferential binding to Rheb-GDP. **a, b** Merged representative confocal micrographs of prey GFP-hMETTL5 (green, *left* panel, control) or GFP-hCIB (*right* panel) with prey HA-GST-Rheb (red) and nanoTRAP in COS-7 cells. Zoomed images of the inset show reverse color images of indicated constructs at the tip of filopodia. * indicate absence at the filopodia tips indicate no interaction of HA-GST-Rheb with the GFP-hCIB2. *Arrowheads* indicate accumulation of indicated construct at the tip of filopodia. Right panel shows interaction of CIB2 with Rheb, the mTORC1 activating GTPase. *Scale bar:* 10 μm. The prey/bait ratio is quantified in **b.** Quantification for nanoTRAP assay. *** *p*< 0.001 by two-way ANOVA, NS – not significant. **c** Strength of CIB2 binding with various Rheb variants from panels **d-f,** and the relative GTP and GDP binding of Rheb variants is estimated from previous studies ^34–36^. NA – data not available. **d** Representative immunoblots of co-IP experiments with indicated HA-GST-Rheb constructs with WT tdTomato-CIB2 show stronger binding with the ‘nucleotide empty’ Rheb-S20N variant. **e** Representative immunoblots of co-IP experiments with indicated HA-GST-Rheb constructs with WT tdTomato-CIB2 show stronger binding with the ‘nucleotide empty’ Rheb-D60K protein and reduced binding with the farnesyl mutant Rheb-C181S. *Short exposure* ~1min, *long exposure* ~10 min. **f** Representative immunoblots of co-IP experiments with indicated HA-GST-Rheb constructs with WT tdTomato-CIB2 show reduced binding with the farnesyl mutants Rheb-C181S and Δ181-184, and cells treated with farnesyl transferase-inhibitor lonafarnib (1000 nM).

CIB2’s preferential binding to nucleotide reduced/empty state of Rheb was further validated by loading glutathione bead-immobilized HA-GST-Rheb (WT) with either GDP or non-cleavable GTPγS and adding CIB2 lysate (**Supplementary Fig. 5b**), and through analyzing additional mutagenic constructs (nucleotide binding capacity summarized in **Fig. 3C**). Previously, a mutagenesis screen and subsequent targeted substitution of aspartic acid at position 60 (D60) to valine (V) or isoleucine (I) showed significantly diminished affinity to GTP but comparable GDP binding capacity as WT Rheb. Meanwhile, the D60K variant turned out to be a nucleotide empty variant ^35,36^. In our co-IP studies, we found that CIB2 preferentially binds to, in order of binding strength, nucleotide empty D60K variant, than D60I and D60V (**Fig. 3e**).

Farnesylation of Rheb is essential for nucleotides loading and activation ^34^. To explore the essentiality of Rheb-farnesylation for CIB2 coupling, we performed co-IP studies using the farnesylation variant C181S, and Rheb with the last four amino acids deleted (Δ181-184), which was shown to be essential for proper loading of nucleotides ^34^. We found significantly reduced CIB2 binding in the absence of Rheb farnesylation (**Fig. 3e, f**). These findings were further confirmed by using the farnesyl transferase inhibitor, lonafarnib (**Fig. 3f**). Finally, as we observed CIB2 interaction both with Rheb and Raptor, which are also known to interact with each other ^37^, we wondered if CIB2 impacts interaction of Rheb with Raptor. However, no obvious effect on the binding of Rheb to Raptor in the presence of CIB2 was observed in co-IP assays, which suggests non-competing binding sites of Raptor and CIB2 with Rheb (**Supplementary Fig. 5c**). Taken together, our studies demonstrated that CIB2 modulates mTORC1 activity, and hence autophagy, by preferentially binding to nucleotide empty/ GDP-Rheb (inactive form).

### mTORC1 is also upregulated in RPE/choroid tissues from patients affected with AMD

Previous studies in mouse models ^16,30,38–40^ and humans samples ^17,39,41^ have shown that both non-canonical and canonical autophagy is essential for RPE function. In humans, AMD is characterized by drusen accumulation in the inner collagenous layer of Bruch’s membrane, RPE vacuolization, and accumulation of lipid deposits within the RPE, Bruch’s membrane, and elsewhere. Aging-associated lipid accumulation is further thought to hinder lipid degradation by phagolysosomes and autolysosomes in RPE, thereby exacerbating accumulation of undigested lipids ^42^, and this has been suggested as an important causative factor in dry AMD. Since our CIB2 deficient mice showed similar age-related RPE pathologies, we reasoned that upregulated mTORC1 signaling might be a contributing factor for the autophagy deficits seen in AMD patients as well. To test this, we evaluated RPE/choroid tissues bio-banked from human subjects affected with dry AMD (n=8) along with age-matched control (n=10) tissues (**Supplemental Table 1**). Consistent with observations made in primary RPE cell cultured from AMD patients ^17^, autophagy was impaired in dry AMD RPE/choroid lysates, as exhibited by accumulation of both p62/SQSTM1 and LC3. Further, dry AMD patients’ lysates exhibited reduced CIB2 expression, and higher levels of phosphorylated, but not of total protein levels, ULK1 and 4E-BP1 (**Fig. 5a, b, Supplementary Fig. 6**), indicating hyperactive mTORC1 signaling. These results demonstrate that, similar to *Cib2* mutant mice, dry AMD patients’ RPE/choroid tissue also has reduced CIB2 levels, hyperactive mTORC1 signaling accompanied by autophagy deficits.

**Figure 5:**
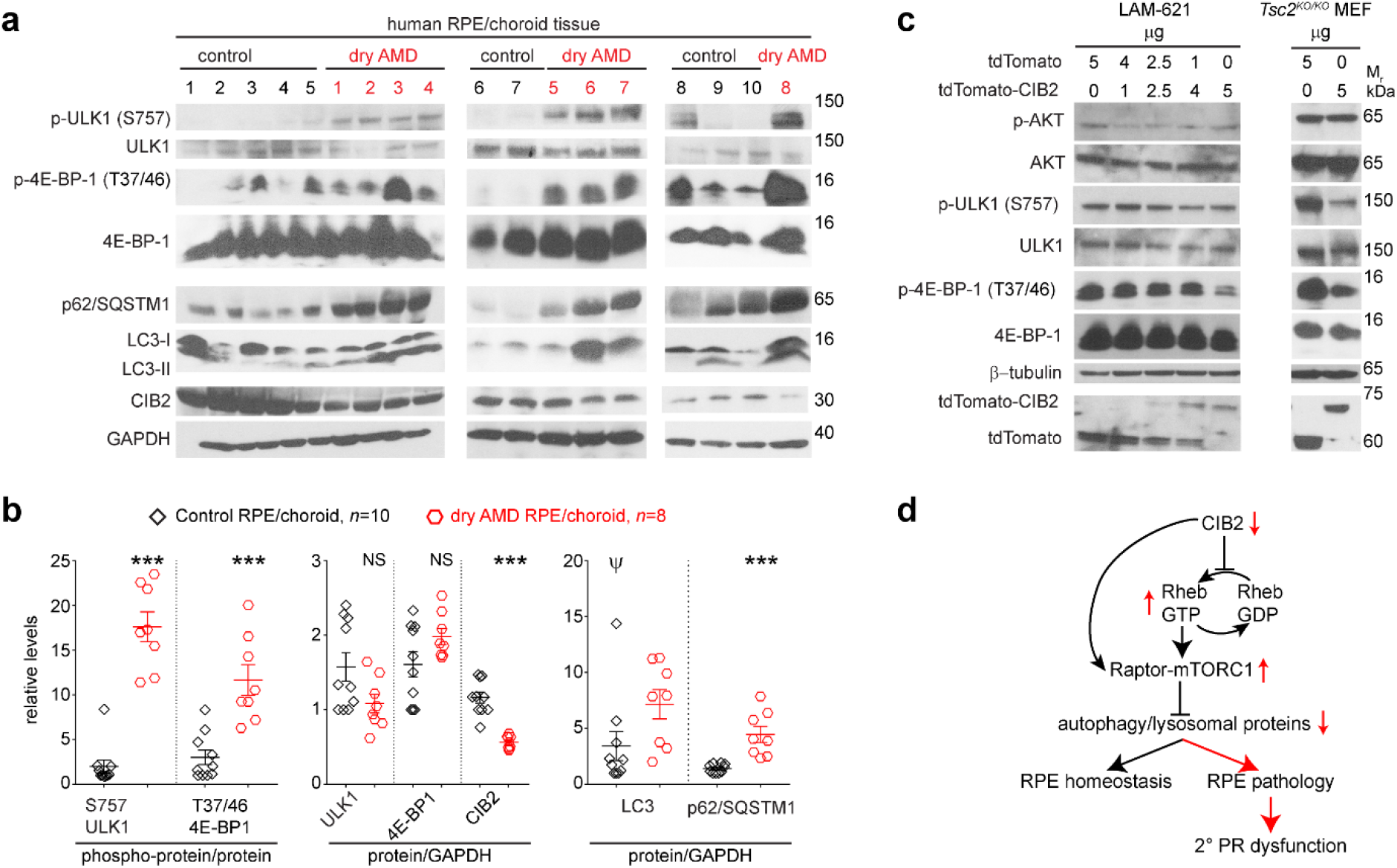
mTORC1 signaling and CIB2 are dysregulated in RPE/choroid lysates from dry AMD cases, while over-expressing CIB2 downregulates mTORC1 signaling in cell lines. **a, b** Immunoblots from RPE/choroid lysates from dry AMD (n=8) or control (n=10) age-matched donors indicate reduced levels of CIB2 and autophagy proteins, and overactive mTORC1 signaling, quantified in panel **b**, concordant to the molecular deficits found in RPE of *Cib2* mutant mice. Unpaired two-tailed *t-*test was used to determine statistical significance. *P-* value of <0.05 (*), <0.01 (**), and <0.001 (***). NS – not significant. Please note that one control sample (*ψ*) is outliner and is highly variant from the rest. Re-quantification, omitting the highly variant control sample, is shown in **supplementary Figure 5**. **c** Immunoblots from lymphangioleiomyomatosis patient-derived cell line (LAM-621; *left* panel) and *Tsc2^KO/KO^*, *p53^-/-^* mouse embryonic fibroblasts (MEF; *right* panel) lysates over-expressing either tdTomato or CIB2. Intriguingly, CIB2 can partially reduce the over-active mTORC1 signaling even in the absence of TSC. **d** Model of CIB2 function in retinal sensory cells. In our model, CIB2 is critical for regulating mTORC1 signaling and autophagy in RPE via its preferential interaction with nucleotide empty or GDP-Rheb (inactive form) and raptor. Loss of CIB2 results in reduced autophagy and exacerbated mTORC1 signaling, thus leading to RPE pathology and secondary photoreceptor (PR) dysfunction.

Besides AMD (shown here), upregulated mTORC1 signaling is also implicated in aging, TSC, and cancers, including lymphangioleiomyomatosis (LAM). Further, TSC variants lead to RPE lesions in human patients ^43^. Above studies demonstrated that CIB2 acts as a negative regulator of mTORC1, and hence, we reasoned that CIB2 overexpression may partially down-regulate the hyperactivated mTORC1 signaling observed in TSC and LAM. To evaluate this, we utilized two different diseases models. First, a patient-derived cell line LAM-621 (LAM-associated renal angiomyolipoma) harboring a variant in TSC2 (p.Arg611Gln) rendering it unable to complex with TSC1, resulting in hyperactivated mTORC1 signaling ^44^. LAM-621 cells overexpressing increasing amounts of CIB2, showed decreasing amounts of phosphorylated 4E-BP-1 and ULK1, but not AKT (**Fig. 5c**, *left* panel). Secondly, *Tsc2^KO/KO^, p53 ^KO/KO^* mouse embryonic fibroblasts (MEFs), similarly overexpressing CIB2, resulted in reduction of mTORC1 but not mTORC2 targets (**Fig. 5c**, *right* panel). Taken together, we show that CIB2 preferentially binds to inactive form of Rheb, thereby regulating mTORC1 activity. Aberrant mTORC1 activity leads to chronic autophagy deficits leading to RPE pathology and secondary photoreceptor dysfunction (**Fig. 5d**). Our findings signify that CIB2 can partially downregulate mTORC1, independently of the TSC complex, and thus potentiating it as a target for modulation of mTORC1 specifically.

## DISCUSSION

This study demonstrates the strength of combined use of mouse models and human disease relevant tissue to uncover functional mechanisms responsible for human disorders. Although, around 35 risk loci for dry AMD have been documented in the literature, the functional mechanism remains elusive, partly due to the fact that often the mouse models do not faithfully recapitulate the full spectrum of human phenotype ^45–48^. Recently, the combination of transcriptome and expression quantitative trait loci (eQTL) analysis on retinal tissues from human AMD patients identified three new genes, *RLBP1, HIC1* and *PARP12.* An association *(p=1.85 x10^-27^*) between a single nucleotide polymorphism (*rs11547207*) in *CIB2* and cis-eQTLs was also documented in that study ^49^. In a parallel study, we reported that both complete loss and haploinsufficiency of CIB2, specifically in RPE, lead to attenuated ERG amplitudes and development of an age-related phenotype encapsulating several dry AMD features ^28^.

Here, we extend the observations and decipher the underlying molecular mechanism of CIB2-mediated regulation of autophagy in RPE. Deficits in autophagy have been associated with age-related pathology in mice and dry AMD in humans ^16,17,30^. Autophagy decreases with ageing in several tissues including the RPE ^19,39,41^. RPE cells cultured from AMD patients showed elevated numbers of autophagosomes, lipid droplets, aberrantly large lysosomes, and higher starvation-induced SQSTM1/p62 levels ^17^. Consistent with these observations, we found an accumulation of both p62/SQSTM1 and LC3, suggesting aberrant autophagy in RPE/choroid samples from dry AMD patients. Concordant with our animal models, we also observed reduced levels of CIB2, and hyperactive mTORC1 in these human samples. Together, these data reconcile CIB2 levels, mTORC1 activity, autophagy, and their age-associated impact on the RPE maintenance and function in mice and humans.

mTORC1, the key negative regulator of autophagy, is cytosolic and in the presence of sufficient nutrients it is recruited to the surface of the lysosomes ^22,23^. GTP-loaded active Rheb allosterically realigns the kinase active site of mTORC1, and thus is a potent and obligate activator of mTORC1 on the surface of the lysosomes ^26^. We found that CIB2 preferentially binds to, in order of binding strength, the nucleotide-empty form of Rheb followed by GDP-loaded inactive Rheb, and then GTP-loaded active Rheb. Generally, for the Ras family of GTPases their respective guanine exchange factors (GEFs) are the ones which bind strongest to the nucleotide empty versions of proteins. However, our current biochemical data do not support the role of CIB2 as a GEF or guanine dissociation inhibitor for Rheb (**supplementary Fig. 6**). Our results here support the function of CIB2 as a “co-factor” to preferentially maintain Rheb in an inactive state. However, further biochemical studies will be needed to decipher the precise mechanism and stoichiometry of CIB2-Rheb and CIB2-Raptor interactions. Furthermore, since mTORC1 is downstream of AMP-activated protein kinase (AMPK), one of the main integrators of energy cues and a negative regulator of mTORC1 ^50^, it would be interesting to assess whether CIB2 established any feedback mechanism between mTORC1 and AMPK pathways.

Besides AMD, a growing literature points to defects in autophagy, lysosomal functions and mTORC1 signaling in ageing and diverse neurodegenerative disorders with complex genetic architecture such as Alzheimer’s, autism, epilepsy, cancers, obesity, and heart diseases. Thus, identifying molecular targets that can be used to modulate or restore autophagy may offer new avenues for treatment of such disorders ^51^. Direct mTOR kinase inhibitors or rapamycin (and its analogs) have shown only limited success in dry AMD patient trials ^52,53^, as chronic exposure to either class of compounds also leads to mTORC2 disassembly and affects cell survival ^54^. Since direct mTOR inhibition is not a viable long term strategy, there is intense efforts to discover drugs modulating the mTORC1 pathway by targeting other players of the pathway such as Rheb ^55^. *CIB2* is a small gene that can easily fit within the carrying capacity of AAV for *in vivo* delivery, supporting potential translatability via gene therapy. Moreover, autophagy is linked to lysosomal biogenesis via the transcription factor EB (TFEB) family, the lysosomal master transcription factor family (TFEB/ TEF3/ MiTF) via which mTORC1 controls lysosomal biogenesis ^56^. The reduction of lysosomal proteins in RPE of *Cib2* mutant mice could potentially be mediated via mTORC1 regulation of TFEB. If so, CIB2 overexpression might be dually beneficial, since it would reduce mTORC1 activity and thereby upregulate autophagy and lysosomal biogenesis - promising topics for further basic or translational studies.

## METHODS

### Animals, tissue harvest, and processing

The ARRIVE and ARVO *Statement for the Use of Animals in Ophthalmic and Vision Research* and the National Institutes of Health *Guide for the Care and Use of Laboratory Animals* guidelines were used for procedures involving and reporting animal research. All animal procedures were reviewed and approved by the IACUC (Institutional Animal Care and Use Committees) of the University of Maryland School of Medicine. Mouse strains are described in the companion study ^28^.

RPE flat mounts were generated by first removing the anterior segment and lens from an eye, and then removing the retina carefully in Hank’s balanced salt solution (HBSS). For autophagy flux assay, whole mounts of RPE/choroid were incubated in DMSO (control) or 50 nM Bafilomycin A1 overnight in DMEM at 37°C, before rinsing with PBS, tap drying on filter paper, followed by snap-freezing on dry ice and storing at 80°C before lysis and immunoblotting. For assessing mTORC1 activity following increased autophagy *in vivo*, mice were fasted with unlimited access to water for 24 hr, followed by RPE/choroid dissection. Eyes were enucleated, opened to remove the lens, retinae dissected from posterior eyecups, and resulting RPE/choroid tissues snap-frozen on dry ice before storing at −80°C before lysis.

### Adipose derived mesenchymal stem cells isolation (MSC), expansion and culturing

MSC were isolated as described ^57^ with some modifications. Briefly, WT and *Cib2^KO/KO^* (n = 3-4/genotype) adipose tissue was dissected from a subcutaneous site of mice at 9 months of age, thoroughly washed with several changes of PBS to remove blood vessels, hairs and other type of connective tissues, and minced. The minced samples were incubated with the collagenase I solution (10 mg solution in DMEM/3 gm tissue) for 40 min at 37°C with shaking. After digestion, sample was diluted 1:1 in culture medium (DMEM supplemented with 15 % FBS and 1xPen-Strep) and filtered to further disintegrate aggregates. The resulting filtrate was centrifuged at 1200 rpm for 10 min, and resulting pellets were re-suspended in 1 ml culture medium and plated the isolated stromal vascular fraction (SVF: consisted of MSC along with circulating blood cells, fibroblasts, pericytes and endothelial cells) in T-25 culture flasks coated with 1% gelatin. WT and mutant MSCs were grown in 6 well plates on cover slip, for amino acid and insulin stimulation experiments, cells were processed as described ^33^.

### Cell culture and nanoSPD

RPE-J cells (ATCC, Manassas, VA) were maintained at 32°C and 8% CO_2_ in 4% FBS/DMEM supplemented with 1x penicillin/streptomycin. HEK-293T, COS-7, LAM-621, and *Tsc2^KO/KO^ p53^KO/KO^* MEF cells were maintained at 37°C/ 5% CO_2_ in 10% FBS-DMEM supplemented with 1x penicillin/streptomycin. before lysis in RIPA buffer with a solution containing 1x protease inhibitors, 1 mM Na orthovanadate, 10 mM Na glycerophosphate, and 10 mM NaF, and stored at −80°C until immunoblotting.

60-70% confluent COS-7 cells in 6-well plates for nanoTRAP were transfected with Lipofectamine 2000 (3:1 ratio) with 1μg plasmid construct each (nanoTRAP, GFP-tagged bait, and prey). 24 hrs after transfection, cells were split 1:10 ratio on glass coverslips to allow for filopodia formation, and fixed 24 hrs later with 4% PFA for 15 min at RT and permeabilized with 0.2 % Triton X-100 in PBS for 15 min at RT, followed by blocking with 10% normal goat serum (NGS) in PBS for at least 30 min at RT. Primary antibodies were diluted in 3% NGS-PBS and incubated overnight at 4°C, followed by the incubations with the indicated goat secondary antibodies. A Zeiss 710 laser scanning confocal microscope or Nikon W1 spinning disk microscope was used for image acquisition, with step size of 0.5 μm. Fiji (ImageJ) ^58^ was used to process images. Finally, LAM-621 and *TSC2^-/-^, p53^-/-^* MEFs were transfected with Lipofectamine 2000 following manufacturer’s instructions.

### Transfection, immunoprecipitation, and immunoblotting

80% confluent HEK293 cells in 10-cm dishes were transfected for co-IP experiments with 10 μg of indicated plasmid/s using the polyethylenimine (PEI) method (3:1::PEI:DNA). 2 plates per condition were used. 36 hrs later the plates were chilled on ice, washed once with ice-cold PBS, and cells collected in ice-cold PBS and centrifuged for 2 min at 1500 RPM at 4°C. Cell pellet was then lysed in IP buffer (0.3% CHAPS, 40 mM HEPES, 2.5 mM MgCl_2_, 1x Protease inhibitors, 1 mM Na orthovanadate, 10 mM Na glycerophosphate, 10 mM NaF) on ice for 20 min, followed by centrifuging at full speed for 20 min at 4°C. Lysates were pre-cleared with control agarose beads for 1 hr rotating at 4°C. 30 μl of a 50% slurry of either HA or GST beads washed 3 times in lysis buffer was added. Samples were rotated 3 hrs or overnight at 4°C, washed 4 times in IP buffer, boiled in 50-100μl SDS sample buffer at 95°C, separated by SDS-gel electrophoresis, transferred to PVDF membranes, and immunoblotted.

For GTPγS or GDP loading experiments, cleared lysates of HEK293 cells over-expressing HA-GST-Rheb were prepared as described in IP buffer lacking MgCl_2_. Rheb was immobilized on GST beads by incubating for 2 hr, rotating at 4°C. Beads were washed 4 times with IP buffer without MgCl_2_. GTPγS or GDP was added to final concertation of 100 μM and 1 mM, respectively. The tubes were incubated with shaking for 1 hr at 37°C. Loading was stopped by placing tubes on ice and adding MgCl_2_ to final concentration of 60 mM. TdTomato (control) or tdTomato-CIB2 cleared lysates were prepared as above (in IP buffer) and dividing equally to GTPγS of GDP loaded Rheb and incubated with rotation overnight. Next day the beads were washed, processed, and immunoblotted as above.

### Nucleotide exchange assays

hCIB2 was subcloned into pET21a (histidine tag on the C-terminus; hCIB2-6xHis) and GST-Rheb in pGEX-4T-2 (Addgene #15889), transformed in Bl2 (DE3) bacteria, cultured in LB at 37°C and induced at OD600 = 0.7-0.8 with 1 mM of isopropyl β-D-1-thiogalactopyranoside at 37°C for ~5 hrs. Pellets were generated by spinning down at 15,000 RPM for 20 mins at 4°C, and frozen at −20°C. For purifying CIB2 protein, cells pellet was lysed in 6M guanidinium lysis buffer provided with the ProBond purification system (ThermoFisher Scientific). The buffer was supplemented with 1mg/ml lysozyme, 5 μg/mL DNase I, protease inhibitor cocktail. Purification was performed using denaturing conditions protocol provided by the manufacturer and elution with denaturing buffer supplemented with 500 mM imidazole. GST-Rheb cell pellet was lysed using IP buffer with 1% CHAPS supplemented with 1mg/ml lysozyme, 5 μg/mL DNase I, protease inhibitor cocktail. GST kit (GE Healthcare, Chicago, IL) was used for purifying GST-Rheb following manufactures instructions and eluted with reduced glutathione. Purified proteins were exchanged in PBS and concentrated with centricon tubes (10k and 30k cutoff respectively).

Rheb at 10 μM was first preloaded with GDP in the nucleotide exchange buffer containing 10 μM GDP (25 mM HEPES, pH 7.4, 100 mM NaCl, 1 mM dithiothreitol, 0.5 mM MgCl_2_, 1 mM EDTA, 10 μM GDP) at 30°C for 2 hrs. An 100 μl reaction containing 1 μM Rheb preloaded with GDP was then assayed for GTPγS binding in the presence or absence of 10 μM CIB2 in the nucleotide exchange buffer containing 5 μM GTPγS (25 mM HEPES, pH 7.4, 100 mM NaCl, 1 mM dithiothreitol, 0.5 mM MgCl_2_, 1 mM EDTA, 5 μM GTPγS, [^35^S]GTPγS for specific activity of ~10,000 cpm/pmol). The reactions were incubated at 30°C and 5 μl of each reaction mixture was taken out at different time points and added into 2 ml of ice-cold TNMD (20 mM Tris, pH8.0, 100 mM NaCl, 10 mM MgCl_2_, and 1 mM dithiothreitol) to terminate the reactions. Protein-bound nucleotide was trapped on nitrocellulose, and the bound radioactivity was counted by liquid scintillation.

### Human AMD/ control tissue

Studies on human tissue followed the Declaration of Helsinki guidelines. Human age-matched RPE-choroid tissues were obtained from donor eyes harvested from local and national eye banks, under an IRB exemption issued by the University of Maryland-Baltimore. A complete ocular history, relevant medical history and age was obtained from each donor. Donors with dry AMD and age-matched normal donors were selected for use. We excluded donors with any eye-related diseases besides AMD, or systemic disorders that might affect retinal/RPE or choroidal function. Eyes were examined and RPE/choroidal tissue isolated by a trained Ophthalmologist (SLB) and any additional lesions noted prior to preservation. Tissues were snap-frozen on dry ice and stored at −80°C prior to use.

### Data analysis and availability

For *in vivo* experiments, at least 3 mouse eyes (from different animals) were used for immunoblots, at least 3 images were averaged per eye from 3 eyes/indicated genotype/time point for confocal microscopy. For cell culture (*in vitro*) experiments, at least three independent experiments or 2 independent experiments for co-IP studies were performed. One-way ANOVA with Tukey’s post-hoc test or Student’s *t*-test was used to compare control sample to test samples, with data presented as mean ± SEM. Differences with p < 0.05 were considered significant. Data were analyzed using GraphPad Prism (GraphPad Software, Inc., La Jolla, CA). Source data are provided with this paper.

| REAGENT or RESOURCE | SOURCE | IDENTIFIER |
| --- | --- | --- |
| **ANTIBODIES (Anti)** |  |  |
| CIB2 (clone 59, homemade) | This paper |  |
| Raptor (clone 24C12) | Cell Signaling Technology | # 2280S |
| mTOR Substrates Antibody Sampler Kit | Cell Signaling Technology | # 9862T |
| TSC1 | Cell Signaling Technology | # 4906S |
| TSC2 | Cell Signaling Technology | # 4308S |
| Actin (clone 13E5) | Cell Signaling Technology | # 4970S |
| β-Tubulin (clone 9F3) | Cell Signaling Technology | # 2128S |
| SQSTM1/p62 (clone D1Q5S) | Cell Signaling Technology | #39749 |
| Phospho-AKT 1/2/3 (ser 473) | Santa Cruz Biotechnology | sc-7985 |
| AKT 1/2/3 | Santa Cruz Biotechnology | sc-8312 |
| ULK1 (clone D8H5) | Cell Signaling Technology | # 8054S |
| Phospho-ULK1 (Ser757) (D7O6U) | Cell Signaling Technology | #14202 |
| GAPDH (clone 0411) | Santa Cruz Biotechnology | sc-47724 |
| HA.11 Epitope Tag | Biolegend | # 901501 |
| AMPK Substrate Antibody Sampler Kit | Cell Signaling Technology | #35277 |
| LC3B | Novus Biologicals | NB100-2220 |
| LAMP1 | Sigma | L1418 |
| Rhodopsin Antibody, clone RET-P1 | EMD Milipore | MAB5316 |
| DsRed (recognizes tdTomato, clone OTI4C8) | OriGene | # TA180084 |
| CIB2 | Abnova | H00010518-A01 |
| **REAGENTS** |  |  |
| Bafilomycin A1 | Sigma-Aldrich | B1793 |
| Rapamycin | Sigma-Aldrich | R0395 |
| Pierce™ Anti-HA Agarose | Thermo Fisher | 26181 |
| Glutathione Sepharose® 4B | GE Healthcare | GE17-0756-01 |
| Pierce™ Control Agarose Resin | Thermo Fisher | INV-26150 |
| ECL™ Prime Western Blotting System | Thermo Fisher | 32106 |
| Chaps Hydrate >=98% (hplc) | Sigma-Aldrich | SIG-C3023-25G |
| Gtp-gamma-s | Abcam | ab146662 |
| Guanosine 5'-diphosphate Sodium Type I | Sigma | G7127 |
| Lipofectamine™ 2000 Transfection Reagent | Thermo Fisher | 11668019 |
| In-Fusion® HD Cloning Plus | Clonetech | 638911 |
| QuikChange Lightning Site-Directed Mutagenesis Kit | Agilent | 210518 |
| Accutase | Sigma-Aldrich | SIG-A6964 |
| ProBond Purification system | Invitrogen | 45-0055 |
| **Experimental Models: Cell Lines** |  |  |
| RPE-J | ATCC | # CRL-2240 |
| HEK-293 | ATCC | # CRL-1573 |
| LAM-621 | Kind gift from Dr. Henske | Yu et al., 2004 |
| *Tsc2*<sup>-/-</sup>, *p53*<sup>-/-</sup> MEF |  | Zhang et al., 2003 |
| **Recombinant DNA** |  |  |
| pRK5-HA-mCherry-raptor | Addgene | 73386 |
| pRK5-HA-GST-Rheb1 | Addgene | 19310 |
| pRK5-myc-mTOR | Addgene | 1861 |
| pRK7-FLAG-TSC1 | Addgene | 8995 |
| pRK7-FLAG-TSC2 | Addgene | 8996 |
| eGFP-C2-hCIB2 | This paper |  |
| tdTomato-C1-mCIB2 | This paper |  |
| ptdTomato Vector-C1 | Clonetech | 632531 |
| pAAV-CIB2-IRES-AcGFP | This paper |  |
| pAAV-AcGFP | Grousbeck Gene Therapy Center |  |
| pcDNA3.1-MYO10-HMM-Nanotrap | Addgene | 87255 |
| eGFP-C2-hFAM92A1 |  | Schrauwen et al, 2018 |
| eGFP-C2-hMETTL5 |  | Richard et al, unpublished |
| Human MCRS1 cDNA ORF Clone in Cloning Vector | Sino Biological US Inc. | HG14816-G |
| **Software and Algorithms** |  |  |
| Prism 6.0 | GraphPad, La Jolla, CA |  |
| Adobe Suite | Adobe, CA |  |
| Fiji | https://fiji.sc/ |  |
| **Mouse strains** |  |  |
| *Cib2<sup>tm1a(EUCOMM)Wtsi</sup>* mice | EUCOMM | EM:05417 |
| VMD2-Cre mice | Gift from Drs Sheldon Miller and Joshua Dunaief | Iacovelli et al., 2011 |
| *Opsin-iCre75* | Jax | Stock No: 015850 |
| ROSA26::FlPe knock in mice | Jax | Stock No: 003946 |

### Contact for reagent and resource sharing

Further information and requests for resources and reagents should be directed to and will be fulfilled by the Lead Contact, Zubair M. Ahmed, PhD.

## Contributions

S.S. and Z.M.A. designed and conceived the project. S.S. carried most of the experiments and analyzed data, X.J. carried out GTP exchange assays. S.R., Sh.R., P.R. and Z.M.A., provided resources, supervised the experiments, and analyzed data. S.L.B. provided human AMD/control tissues, and analyzed data. S.S. and ZMA wrote the manuscript. All authors read, edited, and approved the manuscript.

## Acknowledgments

We thank Ms. D. Gomes, S. Riaz for technical assistance with mouse colony and MSC cultures, respectively, Dr. E. Henske for cell lines, Dr. Marcos Sotomayor for pET21a vector, and the UMSOM core facility for use the Zeiss-710, Nikon W1 microscopes. We thank Drs. T. Friedman, R. Hertzano, C. Mueller, G. Frolenkov, and Ms. M. Ahmed for critically reading the manuscript. This work was supported by NIDCD/NIH grants R01DC012564, R01DC016295 (Z.M.A.), R56DC011803 (S.R.), and partly by the Loris Rich Postdoctoral fellowship, International Retinal Research Foundation, AL, USA (S.S.), and Research to Prevent Blindness Award (Z.M.A.).

